# GABA levels in ventral visual cortex decline with age and are associated with neural distinctiveness

**DOI:** 10.1101/743674

**Authors:** Jordan D. Chamberlain, Holly Gagnon, Poortata Lalwani, Kaitlin E. Cassady, Molly Simmonite, Rachael D. Seidler, Stephan F. Taylor, Daniel H. Weissman, Denise C. Park, Thad A. Polk

## Abstract

Age-related neural dedifferentiation – a decline in the distinctiveness of neural representations in the aging brain–has been associated with age-related declines in cognitive abilities. But why does neural distinctiveness decline with age? Based on prior work in non-human primates and more recent work in humans, we hypothesized that the inhibitory neurotransmitter gamma-aminobutyric acid (GABA) declines with age and is associated with neural dedifferentiation in older adults. To test this hypothesis, we used magnetic resonance spectroscopy (MRS) to measure GABA and functional MRI (fMRI) to measure neural distinctiveness in the ventral visual cortex in a set of older and younger participants. Relative to younger adults, older adults exhibited lower GABA levels and less distinct activation patterns for faces and houses in the ventral visual cortex. Furthermore, individual differences in GABA within older adults positively predicted individual differences in neural distinctiveness. These results provide novel support for the view that age-related reductions of GABA contribute to age-related reductions in neural distinctiveness (i.e., neural dedifferentiation) in the human ventral visual cortex.

## 1. Introduction

Fluid processing abilities (cognitive abilities that do not depend on how much you know) often decline with age, even in the absence of disease (Park et al., 2002). Age-related neural dedifferentiation, that is a decline in the distinctiveness of neural representations with age, has been hypothesized to play a role in such impairments (see Koen & Rugg, 2019 for a recent review). For example, the neural activation patterns associated with different categories of visual stimuli (e.g., faces and houses) are more confusable in older compared to younger adults (Park et al., 2004; Carp et al., 2011) and individual differences in such neural distinctiveness have been found to account for as much as 30% of the variance in fluid processing ability among older adults (Park et al. 2010). Furthermore, computational models have demonstrated that declines in neural distinctiveness can account for a number of age-related behavioral impairments (Li & Lindenberger, 1999; Li, Lindenberger, & Sikström, 2001; Li & Sikström, 2002). Conversely, neural distinctiveness is not associated with crystallized intelligence (cognitive processing that depends critically on knowledge), which typically does not decline with age.

An important open question is why neural distinctiveness declines with age. One hypothesis is that age-related reductions in the brain’s major inhibitory neurotransmitter, gamma aminobutyric acid (GABA), play a role (Hua et al., 2008). GABA is critical for resolving cortical competition between different representations (Isaacson & Scanziani, 2011), so a reduction in GABA among older adults might plausibly lead to less distinct activation patterns for competing stimuli. Furthermore, some evidence suggests that GABA levels in visual cortex are reduced in older compared to younger adults (Simmonite et al., 2018, but see Aufhaus et al., 2013 and Pitchaimuthu et al., 2017) and individual differences in GABA levels in pre-supplementary motor area and occipital cortex predict individual differences in cognitive performance in older adults (Hermans et al., 2018; Simmonite et al., 2018).

Recent work from our group has also found evidence for a link between GABA levels and neural dedifferentiation in humans. Specifically, Lalwani et al. (2019) showed that GABA levels in auditory cortex were significantly associated with the distinctiveness of fMRI activation patterns associated with music and speech. Additionally, Cassady et al. (2018, 2020) found that the segregation of resting state motor networks was positively associated with sensorimotor GABA levels in healthy older adults, and the degree of segregation was associated with individual differences in motor performance.

Findings from the visual cortex of non-human primates further support this hypothesis. Leventhal et al. (2003) found that visual neurons in young monkeys responded more selectively to stimuli with specific orientations and directions than did visual neurons in old monkeys. However, electrophoretic application of GABA, or the GABA_A_ receptor agonist muscimol, made the neurons in older monkeys respond selectively, like the neurons in young monkeys. Conversely, application of the GABA_A_ receptor antagonist bicuculline made visual neurons in young monkeys respond less selectively, like the neurons in old monkeys. Together, these results demonstrate that changes in GABA activity can cause changes in neural selectivity at the level of individual receptive fields in non-human primates. Whether age-related reductions of GABA activity play a role in age-related reductions of neural distinctiveness in human visual cortex remains unexplored.

Inspired by these prior findings, we tested this hypothesis. Specifically, we combined magnetic resonance spectroscopy (MRS) with functional MRI (fMRI) to measure both GABA levels and neural distinctiveness in the ventral visual cortex of young and old adults. We hypothesized that both measures would be reduced in the older group compared with the younger group. We also hypothesized that older adults with higher levels of GABA would also exhibit the most neural distinctiveness.

## 2. Materials and Methods

### 2.1 Ethics Statement

The University of Michigan Institutional Review Board approved the study procedures. All participants provided verbal and written consent prior to the study.

### 2.2 Participants

37 younger adults (mean age = 23 years, SD = 3, range = 18-29, female = 20) and 51 older adults (mean age = 70 years, SD = 5, range = 65-87, female = 31) from the local Ann Arbor community participated in the experiment as part of the Michigan Neural Distinctiveness (MiND) project (Gagnon et al., 2019). All participants were native English speakers, right-handed, physically and psychologically healthy, not taking vascular or psychotropic medication, had normal or corrected-to-normal vision, and had no other MRI contraindications. Finally, all participants scored greater than 26 on the Montreal Cognitive Assessment (MoCA; Nasreddine et al., 2005). See Gagnon et al. (2019) for more details regarding the MiND project design.

### 2.3 Experimental design

Prior to fMRI scanning, participants completed a brief visual acuity test derived from the NIH Toolbox for Assessment of Neurological and Behavioral Function (Gershon et al., 2010). We used this measure to test whether age-related declines in neural distinctiveness might be related to individual differences in peripheral vision. Briefly, participants viewed letters sequentially presented on an iPad at a distance of three meters. Participants verbally stated what letters they saw on the screen, and the letters became smaller after each correct response. The NIH Toolbox software calculates a LogMAR score (modified Snellen visual acuity score) and then converts it to a standard score of visual acuity.

During fMRI scanning, participants completed a visual face/house task similar to that used by Park et al. (2010). The task consisted of one six-minute fMRI run with six 20-second blocks of faces and six 20-second blocks of houses, in pseudorandom order. A 10-second fixation block followed each face and house block. During the face blocks, participants viewed greyscale images of male faces. During the house blocks, participants viewed greyscale images of houses. Each stimulus appeared for 500 milliseconds, after which there was a 500 millisecond interstimulus interval (ISI). We instructed participants to press a button with their right index finger whenever they saw a female face during the face blocks, and whenever they saw an apartment building during the house blocks. Such target images occurred approximately once per minute. We presented the stimuli using E-Prime 2.0 on a back-projection system. We recorded participants’ responses using a Lumina response pad (Cedrus).

### 2.4 fMRI data acquisition

We collected fMRI data with a 3T GE MRI scanner at the University of Michigan’s Functional Magnetic Resonance Imaging Laboratory. We acquired blood oxygen level dependent (BOLD) images using a single-shot gradient echo (GRE) reverse spiral pulse sequence (TR = 2000ms, TE = 30ms, FOV = 220mm, voxel size = 3.4375 x 3.4375 x 3mm, 43 axial slices). We collected high-resolution T1 images using an SPGR (3D BRAVO) sequence with the following parameters: Inversion Time (TI) = 500 ms; flip angle = 15°; Field of View (FOV) = 256 x 256 mm; 1 x 1 x 1 mm voxels; 156 axial slices.

### 2.5 Magnetic resonance spectroscopy data acquisition

We collected MRS data using the same scanner during a different scanning session. We placed 3×3×3cm voxels in the ventral visual cortex near the left and right mid-fusiform gyrus (See Figure 1 for overlap in voxel placement in ventral visual cortex across participants). We customized voxel placement for each participant to maximize overlap with ventral visual fMRI activity during the face/house task from the previous fMRI session (using a contrast of Face vs. Fixation and House vs. Fixation). We also collected MRS data from voxels placed in the auditory cortex and sensorimotor cortex and used those data to test whether the relationship between GABA and neural distinctiveness was regionally specific. Auditory cortex spectroscopy voxels were placed to maximize overlap with fMRI activation from a contrast of Speech vs. Fixation and Music vs. Fixation, and sensorimotor cortex voxels were placed to maximize overlap with Left Hand Tapping vs. Fixation and Right Hand Tapping vs. Fixation (Gagnon et al., 2019). We acquired GABA-edited MR spectra using a MEGA-PRESS sequence using the following acquisition parameters: TE = 68 ms (TE1 = 15ms, TE2 = 53 ms); TR = 1.8s; 256 transients (128 ON interleaved with 128 OFF) of 4,096 data points; spectral width = 5kHz, frequency selective editing pulses (14ms) applied at 1.9 ppm (ON) and 7.56 (OFF); total scan time, approximately 8.5 min per voxel.

**Figure 1.**
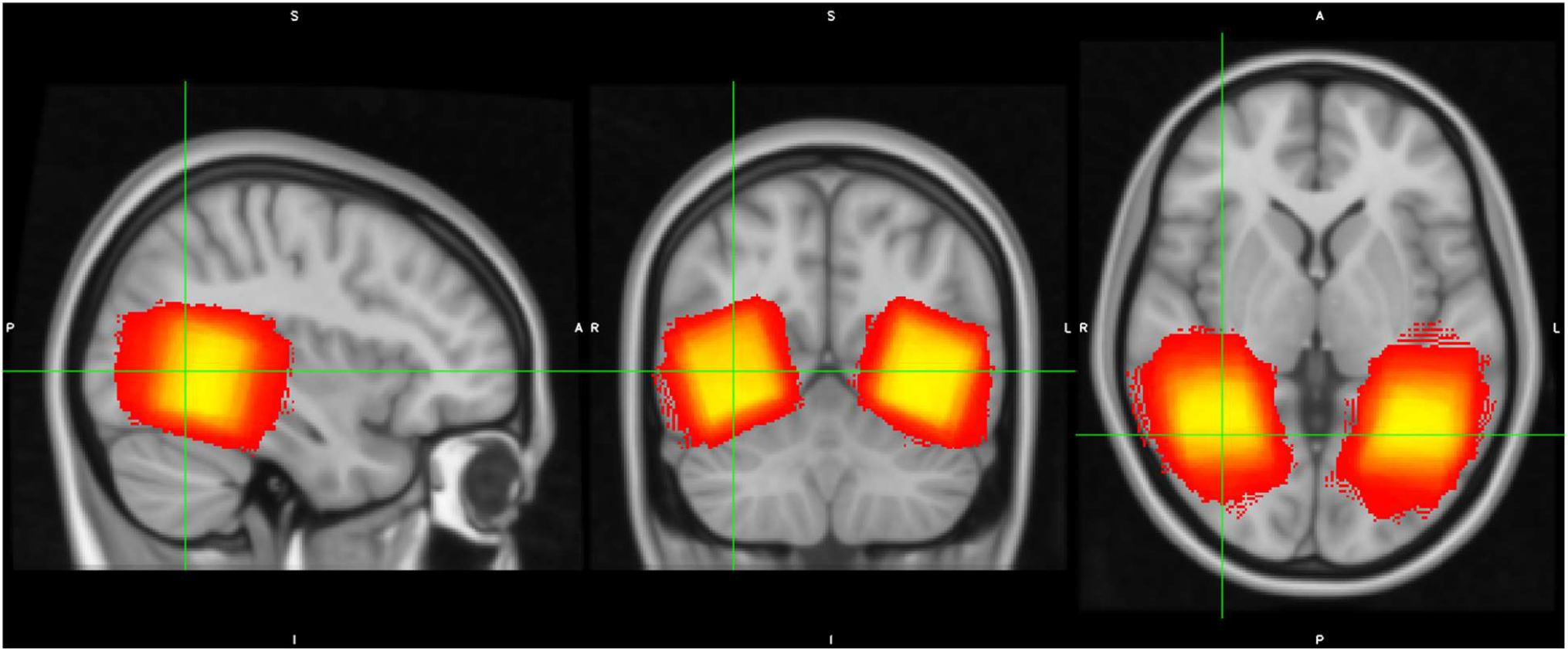
MRS voxel placement. Voxel placement in ventral visual cortex across all participants is depicted on the MNI T1 1mm template. Brighter colors (yellow) indicate greater overlap and darker colors (red) indicate less overlap across participants.

### 2.6 Statistical analysis

#### 2.6.1 fMRI pre-processing

We k-space despiked, reconstructed, and corrected the MRI data for heart beat and respiration using the RETROICOR algorithm (Brooks et al., 2008). We then slice time corrected the data using the spm_slice_timing function from SPM (https://www.fil.ion.ucl.ac.uk/spm) and motion corrected using mcflirt from FSL 5.0.7 (www.fmrib.ox.ac.uk/fsl). We resampled the data into two-dimensional cortical surfaces (one for the left hemisphere and one for the right hemisphere) based on a white/gray matter segmentation of each subject’s own high-resolution structural image computed using Freesurfer’s recon-all function (version 6, https://surfer.nmr.mgh.harvard.edu/). We spatially smoothed the data within each cortical surface using a 5-mm two-dimensional smoothing kernel and used Freesurfer’s FsFast processing stream to fit a general linear model to the fMRI time series at each point, or vertex, on the cortical surface. The model included box-car regressors corresponding to the face and house conditions, convolved with FsFast’s hemodynamic response function.

#### 2.6.2 ROI mask creation

We created participant-specific structural masks of the bilateral fusiform gyrus and bilateral parahippocampal gyrus using the cortical parcellation of each participant’s brain generated by Freesurfer’s recon-all function (this function also generated gray matter volume estimates of these gyri which we used as nuisance covariates in some analyses). We then computed participant-specific functional regions-of-interests (ROIs) within these structural masks based on activation (i.e., beta values) during the face and house conditions. Specifically, we sorted the beta values for each condition at all the vertices within the structural mask from largest to smallest. We added the most activated vertex from both the face and house conditions to the ROI, followed by the second most activated vertex in each condition, and then the third most active, and the fourth, and so on while maintaining an equal number of vertices defined from the face and house conditions. If the next most active vertex in one of the conditions was already included in the ROI (based on the other condition), then we added the next most active vertex from that condition that was not already in the ROI. In the end, this process produced functional ROIs that only included the most activated vertices, and that included an equal number of face-active and house-active vertices. This allowed us to compute the neural distinctiveness of the two activation patterns in an unbiased manner.

Note also that this ROI definition procedure was orthogonal to the hypotheses examining age and GABA levels. The ROI was based on two within-subject comparisons: house vs fixation, and face vs fixation. The most activated vertices from each of these contrasts were included in a given participant’s functional ROI. Neural distinctiveness for faces and houses was then calculated within this functional ROI. However, the hypotheses of interest involved between-subject contrasts: the first examining potential distinctiveness differences between younger and older adults, and the second examining covariance between distinctiveness and GABA levels. The ROI selection procedure was therefore orthogonal to the hypotheses being tested.

We used this process to create functional ROIs of varying size (from 1,000 vertices up to the size of the entire structural mask), which allowed us to test whether any observed age-related differences in neural distinctiveness depended on the number of vertices selected. See Figure 2 for a participant-specific example with a functional ROI of 5,000 vertices.

**Figure 2.**
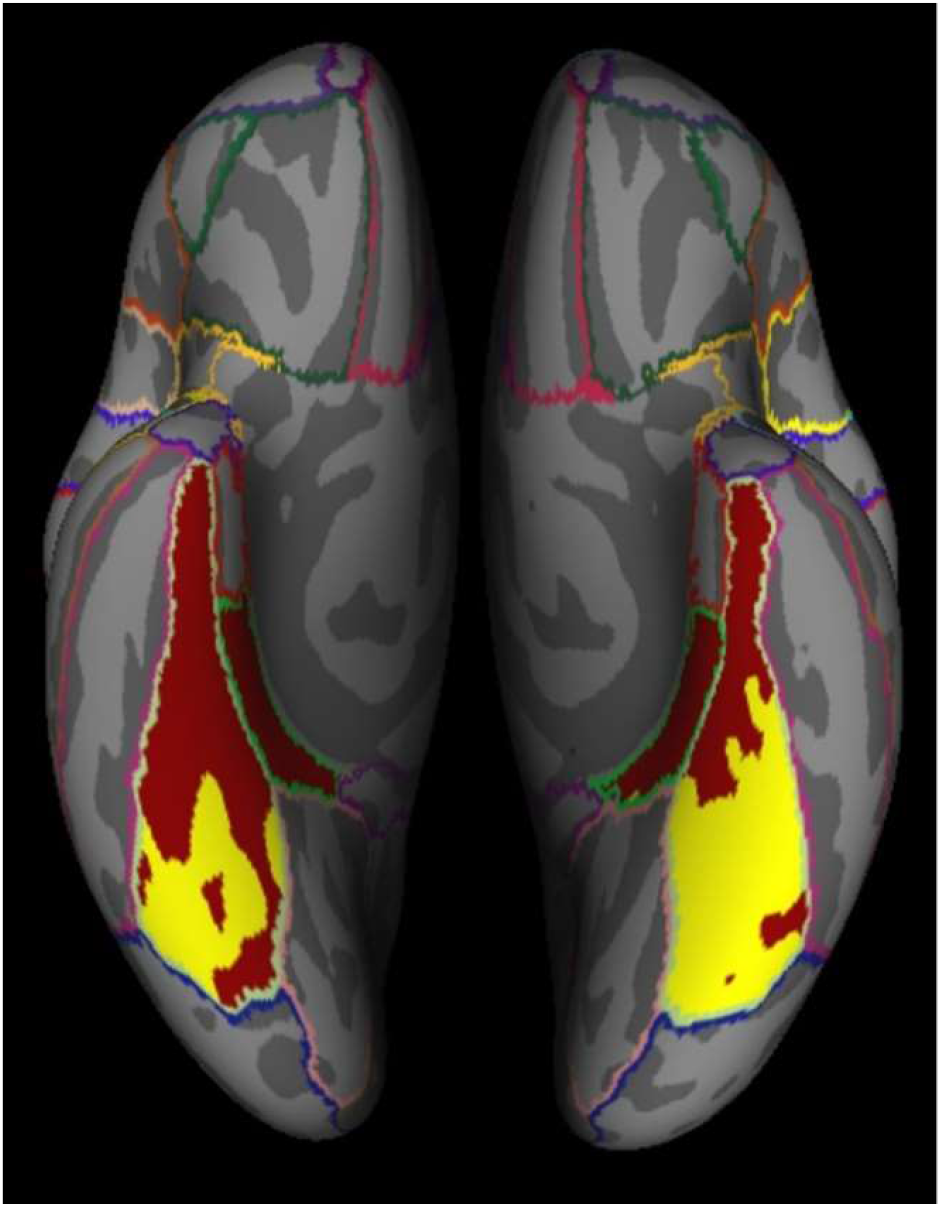
Participant-specific example of ventral visual structural (red) and functional (yellow) ROIs. The functional ROI was created using the 5,000 most activated vertices during the face and house conditions.

#### 2.6.3 Neural distinctiveness

Following Haxby et al. (2000) and previous work by Park et al. (2011), we computed the distinctiveness of face and house activation patterns by subtracting (1) the similarity of activation patterns from different conditions (i.e., faces versus houses) from (2) the similarity of activation patterns within the same condition (i.e., faces versus faces and houses versus houses). If face and house activation patterns are highly distinct, then within-condition similarity should be greater than between-condition similarity. On the other hand, if face and house patterns are less distinct, then within- and between-condition similarity should differ to a lesser degree. While the accuracy of a machine learning classifier (e.g. a support vector machine) is often used as a measure of neural distinctiveness (Park et al., 2010; Fandakova et al., 2019), our use of a block design with 12 blocks meant that classification accuracy could take on only one of 13 discrete values corresponding to the number of 12 blocks that were correctly classified (e.g 0/12, 1/12, 2/12 etc). Furthermore, classifier accuracy was at ceiling for the majority of participants. We therefore used the correlation-based measure as it provides a more fine-grained, continuous measure of distinctiveness that is not susceptible to ceiling effects (Park et al., 2011).

We computed within- and between-condition similarity by calculating (1) the average (Fisher-transformed) correlation between all pairs of blocks within the same condition (i.e., all the face-face pairs and house-house pairs) and (2) the average (Fisher-transformed) correlation between all pairs of blocks from different conditions (i.e., all the face-house pairs). We defined neural distinctiveness as the difference between these measures of within- and between-condition similarity. As this measure of neural distinctiveness is the difference between two correlations, it can range from -2 to 2 with higher numbers indicating greater distinctiveness and lower numbers indicating less distinctiveness.

To assess age differences in neural distinctiveness we used pairwise t-tests separately at each functional mask size of 1,000, 2,000, 5,000, and 10,000 vertices, as well as the full anatomical mask (∼12,000 vertices). To examine age differences in neural distinctiveness in the context of gray matter volume and visual acuity, we used a partial correlation analysis to examine the effect of age group (younger, older) on neural distinctiveness while controlling for gray matter volume (Voss et al., 2008), visual acuity, and univariate functional activation (face + house vs. fixation). We also examined potential age differences in within- and between-condition similarity separately (Carp et al., 2011). All statistical analyses of neural distinctiveness were conducted using *R* software (R Core Team, 2017).

#### 2.6.4 Magnetic resonance spectroscopy analysis

We used the Gannet 3.0 MATLAB toolbox (Edden et al., 2013) to analyze the MR spectra and estimate GABA levels in each voxel. Gannet performs time domain frequency and phase correction of the MR spectra using spectral registration. The time domain was filtered with a 3-Hz exponential line broadening and zero-filled by a factor of 16. Gannet models the GABA peak using a five-parameter Gaussian model between 2.19 and 3.55 pm, and models the water peak using a Gaussian-Lorentzian. Metabolite concentration values are scaled to water. Gannet also coregisters each MRS voxel to the T1-weighted SPGR image and calls SPM to segment the T1-weighted image. It then uses the results to estimate the fraction of gray matter, white matter, and cerebrospinal fluid within the MRS voxel and computes an estimate of GABA that is corrected for tissue composition, as well as the water relaxation times in CSF, white matter, and gray matter (equation number four in Harris et al., 2015). Finally, we note that the GABA peak in the GABA-edited spectrum is contaminated by coedited macromolecules. In keeping with previous reports (Edden et al., 2014), we will refer to our measurements using the term GABA+ (i.e., GABA + macromolecules) to reflect this limitation.

We first used planned t tests to examine age differences in tissue-corrected GABA+ levels. We then assessed age differences in GABA+ levels controlling for differences in signal quality (SNR, fit error, and FWHM). We used a 2 (age group: younger, older) x 2 (hemisphere: left, right) mixed design ANOVA (age group as a between-subjects factor and hemisphere of voxel placement as a within-subjects factor) and GABA+ levels as the dependent measure. To examine individual differences in neural distinctiveness and tissue-corrected GABA+ within older adults, we conducted simple linear regression using tissue-corrected GABA+ as our predictor variable and neural distinctiveness as our outcome variable. To assess whether the relationship between neural distinctiveness and GABA+ was specific to the ventral visual cortex, we conducted a multiple linear regression using tissue-corrected sensorimotor cortex and auditory cortex GABA+ estimates as predictor variables and ventral visual neural distinctiveness as the outcome variable while controlling for age and gray matter volume. We also repeated the above statistical tests using raw GABA+ estimates instead of tissue-corrected GABA+, reported in supplemental materials. All statistical analyses of GABA+ were conducted using *R* software (R Core Team, 2017) and significance was defined at the *p* = .05 level.

## 3.0 Results

### 3.1 GABA+ level

First, we assessed potential age differences in GABA+ levels in the ventral visual cortex. Average tissue-corrected GABA+ levels were significantly lower in the old than in the young adults (*t*(84.59) =4.67, *p* < .001; Figure 3), We also observed a significant age-related reduction in SNR (*t*(82.70) =-2.41, *p* < .05), but not in fit error or FWHM (both *p*’s > .05) in Gannet’s fits to the GABA peaks. Age differences in tissue-corrected GABA+ levels remained significant after controlling for SNR differences (r(87) = -0.38, *p* < .001). Left and right hemisphere GABA+ measures were significantly correlated when using tissue-corrected GABA+ (r(84) = .46, *p* < .001), and a mixed design ANOVA (age group as a between-subjects factor and hemisphere of voxel placement as a within-subjects factor) found no significant interaction between hemisphere and age (*F*(1,84) = .29, *p* > .05). We therefore averaged the GABA+ measures from the two hemispheres for subsequent regression analyses (*see 3*.*3 Individual differences in neural distinctiveness and GABA+ levels*). We repeated the above tests using raw GABA+ levels, rather than tissue-corrected levels, and found no differences in outcomes (see Supplemental Materials).

**Figure 3.**
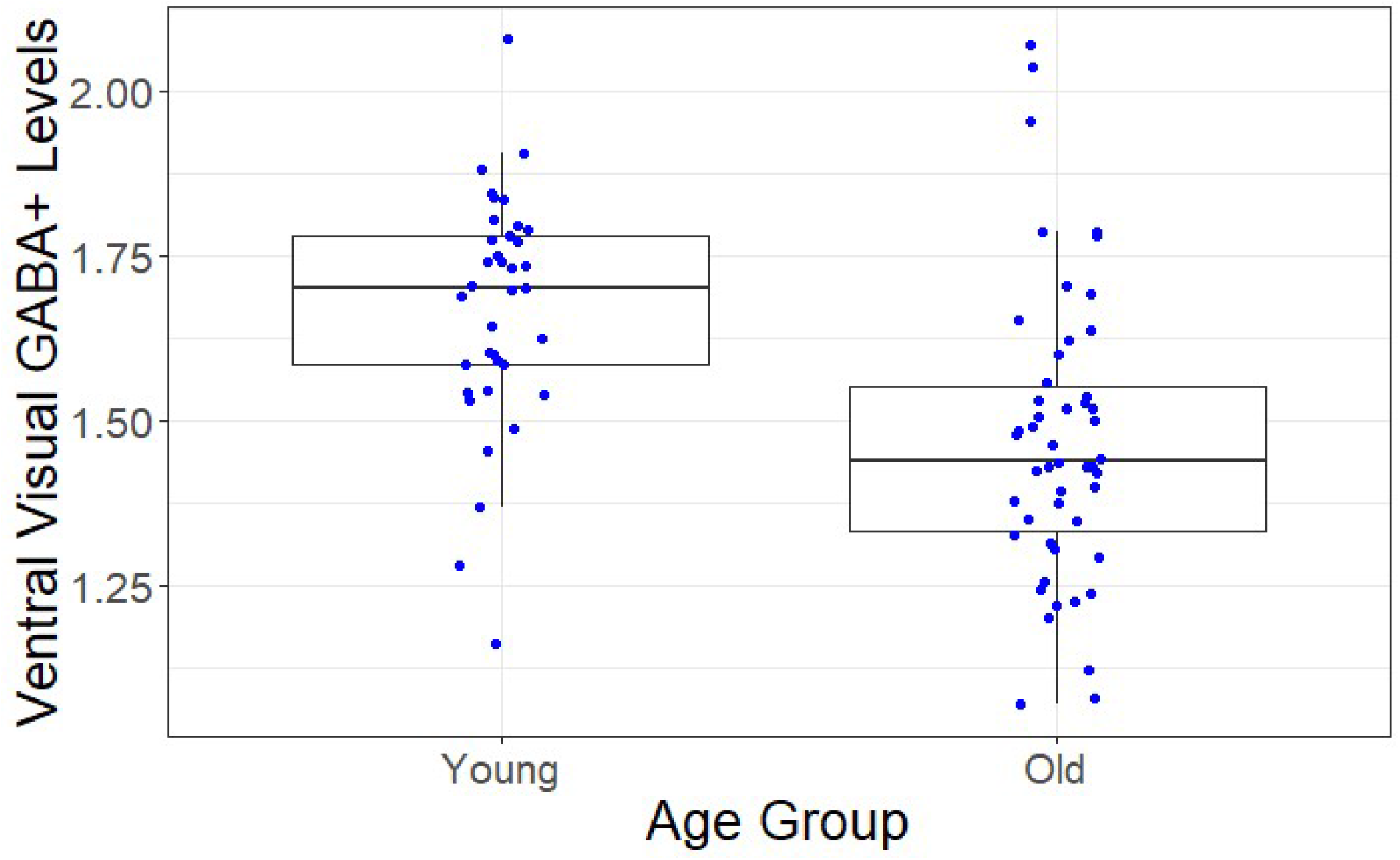
Age-related differences in ventral visual cortex GABA+ levels. GABA+ was significantly reduced in older compared to younger adults.

### 3.2 Neural distinctiveness

Next, we investigated the relationship between age and neural distinctiveness in the ventral visual cortex. Figure 4 plots average neural distinctiveness in young and old adults as a function of ROI mask size. Using pairwise t tests for each functional mask size we found that neural distinctiveness was lower in older adults compared to younger adults across all mask sizes (all *p*’s < .01). As no interaction was present between ROI mask size and age group, we chose an intermediate ROI size of 5,000 vertices for all subsequent statistical analyses using neural distinctiveness. Previous work suggests that age-related declines in regional gray matter volume may contribute to differences in neural distinctiveness (Park et al., 2012) and gray matter volume was lower in the older adults than in the young adults in the fusiform gyrus (left fusiform *t*(70.17) = 4.71, *p* < 0.001; right fusiform *t*(57.76) = 6.16, *p* < 0.001) and parahippocampal gyrus (left parahippocampal *t*(52.82) = 4.65, *p* < 0.001; right parahippocampal gyrus (*t*(69.56) = 5.63, *p* < 0.001). Nevertheless, neural distinctiveness was lower in old adults than in young adults even after controlling for gray matter volume (r(87) = -0.24, *p* < .05).

**Figure 4.**
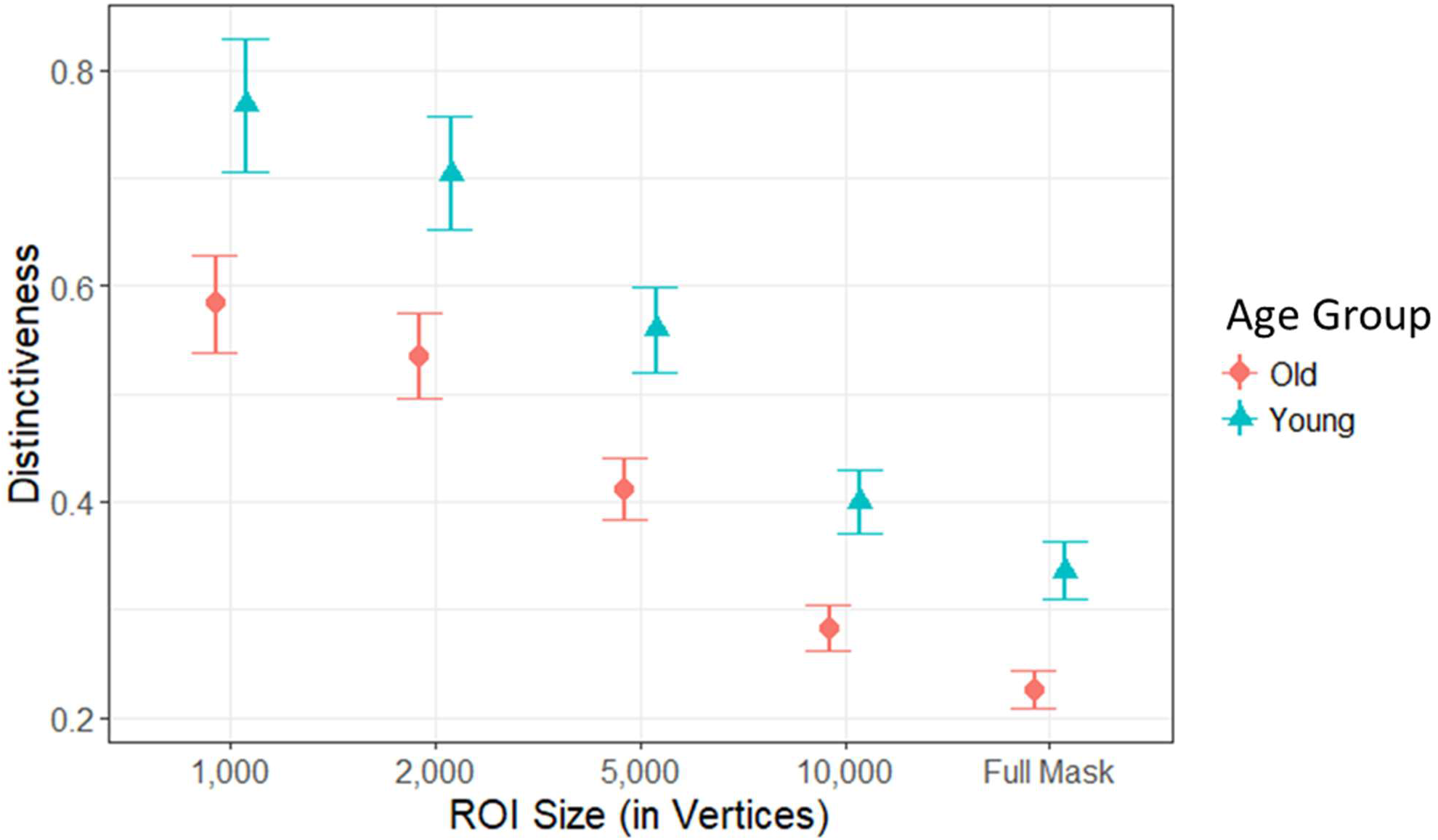
Age-related differences in neural distinctiveness. Neural distinctiveness was significantly reduced within the ventral visual cortex of older (orange circles) compared to younger adults (green triangles) across a range of mask sizes (from 1,000 vertices up to the Full Anatomical Mask).

It is also possible that age-related declines in peripheral visual ability (Greene & Madden, 1987) contribute to neural dedifferentiation in the visual cortex. However, the older adults in our study (who were allowed to use eyeglasses and/or contacts) did not have significantly reduced visual acuity relative to the younger adults (*t*(81.26) = 0.41, *p* > 0.05), and neural distinctiveness remained lower in the old than in the young adults after controlling for visual acuity (r(87) = - 0.31, *p* < .01).

We also tested whether differences in univariate activity could account for age-related reductions in neural distinctiveness. Older adults in our sample did exhibit reduced univariate activity in the face + house vs. fixation contrast (*t*(84.08) = -2.31, *p* < 0.05), but we still observed reduced neural distinctiveness in older compared to younger adults after controlling for these differences (r(87) = -0.26, *p* < .05).

In a previous study, we found evidence for both an age-related increase in between-condition similarity and an age-related decrease in within-condition similarity (Carp et al., 2011). In the present study, however, we only found evidence of an age-related decrease in within-condition similarity (*t*(80.64) = -3.90, *p* < .0001) without a significant age-related change in between-condition similarity (*t*(70.72) = - 0.38, *p* > .05).

### 3.3 Individual differences in neural distinctiveness and GABA+ levels

We also examined the relationship between individual differences in GABA+ and individual differences in neural distinctiveness within older adults through multiple linear regression. Older adults with higher levels of tissue-corrected GABA+ exhibited increased neural distinctiveness (*β*(47) = .27, *p* < 0.05), even after controlling for age, gray matter volume, and GABA+ levels in auditory and sensorimotor voxels within the older adults (*β*(45) = .37, *p* < 0.01; Figure 5). Furthermore, tissue-corrected GABA+ levels in both sensorimotor cortex and auditory cortex failed to predict neural distinctiveness (sensorimotor *β*(45) = -0.06, *p* > 0.05; auditory *β*(45) = -0.26, *p* > 0.05). We also examined the relationship between tissue-corrected ventral visual GABA+ levels and within- and between-condition similarity in older adults. Tissue-corrected GABA+ levels did not significantly predict within- (*β*(49) = .07, *p* > 0.05) or between-condition similarity (*β*(49) = - 0.19, *p* > 0.05). We replicated the above results using raw GABA+ levels rather than tissue-corrected GABA+ levels (see Supplemental Materials. We also replicated the results when using a circular functional ROI of 2,000 vertices centered around each subject’s face + house > fixation activation peak (following the procedures of Park et al., 2012, see Supplemental Materials). Please see Table 1 for descriptive summaries for variables of interest.

**Figure 5.**
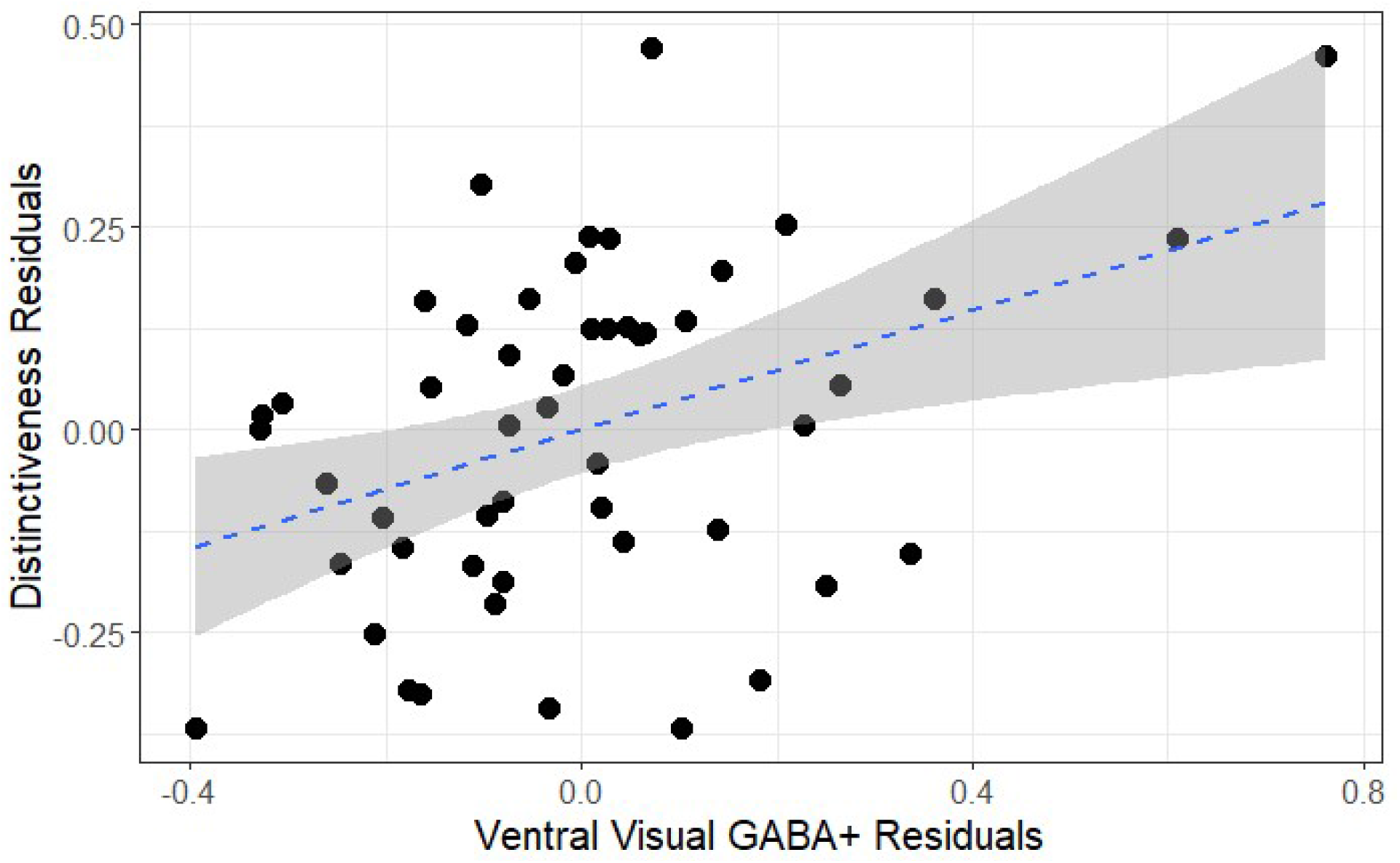
Average GABA+ levels and neural distinctiveness. GABA+ was positively associated with neural distinctiveness in older adults after controlling for gray matter volume, age, sensorimotor GABA+, and auditory GABA+.

**Table 1.** Summary of Key Age Effects.

| Variable | Young Adult | Old Adult |
| --- | --- | --- |
|  | *M (SD)* | *M (SD)* |
| Tissue-corrected Visual GABA+ | 1.83 (.19) | 1.62 (.24) |
| Tissue-corrected Auditory GABA+ | 2.15 (.21) | 1.97 (.24) |
| Tissue-corrected Somatomotor GABA+ | 2.35 (.19) | 2.18 (.20) |
| Neural Distinctiveness | .55 (.24) | .41 (.21) |
| Between-condition Similarity (r) | .07 (.20) | .06 (.17) |
| Within-condition Similarity (r) | .63 (.19) | .47 (.20) |
| Univariate Face Estimate (beta) | .67 (.20) | .50 (.24) |
| Univariate House Estimate (beta) | .22 (.18) | .21 (.19) |

## 4.0 Discussion

*Overview* In this study, we report three main findings. First, GABA+ levels in ventral visual regions were significantly lower in older adults than in young adults. Second, the activation patterns evoked by faces and houses were significantly less distinct in older adults than in young adults. Third, older adults with higher GABA+ levels in the ventral visual cortex exhibited greater neural distinctiveness than older participants with lower GABA+ levels. We discuss each finding in turn.

### 4.2 GABA+ levels decline with age

We found that GABA+ levels in ventral visual cortex were significantly reduced in the older group compared with the younger group. This result extends previous MRS studies that have found age-related declines in GABA+ in frontal and parietal cortex (Gao et al., 2013, Porges et al., 2017, Grachev & Apkarian, 2001) and with a recent meta-analysis suggesting GABA+ levels increase during adolescence and decline throughout later adulthood (Porges et al., 2020). It is also consistent with animal work showing a decline in the number of GABA-immunoreactive neurons in the inferior colliculus (Caspary, Milbrandt, & Helfert, 1995), striate visual cortex (Hua et al., 2008), and hippocampus (Stanley & Shetty, 2004), and with the decline of GABA receptor subunits within primary visual cortex in later adulthood (Pinto et al., 2010). Our results lend support to the notion that GABAergic functioning is reduced in older adults as the visual cortex ages.

On the other hand, our MRS findings differ from those recently reported by Pitchaimuthu et al. (2017), who found increased GABA levels in the visual cortex of old compared to young adults within the banks of the calcarine sulcus. Their MRS voxel placement was significantly more posterior than that used for the voxels in the present study, so one possibility is that GABA+ levels do not change uniformly within the visual cortex in later adulthood. However, we recently collected MRS data from 20 older and 19 young adults in an early visual cortex voxel similar to that used by Pitchaimuthu et al. (2017), and we still found a significant reduction in GABA+ in the older group (Simmonite et al., 2018). Thus, further research will be necessary to resolve this discrepancy.

What might cause the age-related reductions in GABA levels that we observed? Many aspects of GABAergic functioning change with age and could potentially play a role. For example, the density of GABA-immunoreactive neurons in visual cortex is reduced in older vs. younger cats (Hua et al., 2008) and similar results have been reported in other species and brain areas. Stanley et al. (2012) reported an age-related loss of GABAergic interneurons in in the hippocampus of older rats and Majdi et al. (2007) reported an age-related reduction in the density of GABAergic boutons in rat frontal and parietal cortex. Likewise, the number of parvalbumin-containing and somatostatin-containing interneurons has been reported to be reduced in the sensory and motor cortex of older vs. younger rats (Miettinen et al., 1993) and Cha et al. (1997) reported a similar reduction in the number of vasoactive intestinal polypeptide-containing interneurons. Other studies have reported age-related reductions in glutamate decarboxylase, from which GABA is synthesized (e.g., Liao et al. 2016).

Further evidence for age-related impairments in GABA function come from transcranial magnetic stimulation (TMS) studies. Preceding a TMS pulse by a conditioning pulse can reduce the size of the response elicited by the TMS pulse, and this kind of short-interval cortical inhibition (SICI) is mediated by GABA (Ziemann et al., 1996). The original studies of SICI were done in motor cortex, but our group has found a similar SICI-like phenomenon in visual cortex in which the size of visual phosphenes elicited by a TMS pulse is reduced by a preceding conditioning pulse (Khammash et al., 2019). We have also found that this effect is reduced in older adults (unpublished), providing further evidence of GABAergic dysfunction in older adults.

### 4.3 Neural distinctiveness declines with age

In keeping with previous reports of age-related neural dedifferentiation (Park et al., 2004; Payer et al., 2006; Park et al., 2010; Carp et al., 2011; Koen et al., 2019), we found that neural distinctiveness in ventral visual cortex was reduced in older adults relative to younger adults. This effect was present even after controlling for visual acuity, suggesting that age-related declines in peripheral sensory capabilities cannot completely explain declines in neural distinctiveness. We also found that age-related declines in neural distinctiveness were still present after controlling for gray matter volume. This result is consistent with a report from Voss et al. (2008) who found that neural dedifferentiation was not related to local gray matter volume differences in the visual cortex of older adults. Likewise, age-related neural dedifferentiation in motor cortex was reported to survive corrections for local gray matter volume (Carp et al., 2011). Together, these reports suggest that local gray matter differences within the visual cortex cannot completely explain reduced neural distinctiveness.

Neural distinctiveness could be reduced in older compared to younger adults for at least three reasons. The most obvious is that the activation patterns in response to faces and houses are more similar to each other in the older vs. younger participants (i.e., the neural representations for the two categories overlap to a greater degree) (Haxby et al., 2001). If so, then the between-condition similarity measure would increase in older adults and neural distinctiveness (calculated as within-condition similarity minus between-condition similarity) would decrease.

A second possibility is that the neural representations *within* each stimulus category become noisier and less reliable. For example, if the activation evoked by faces is inconsistent from block to block then the within-condition similarity of the face blocks will decline, and so will neural distinctiveness. This could result from reduced representational fidelity of individual stimuli (St-Laurent et al., 2014; Zheng et al., 2018), as has been shown within blocks of faces (Goh et al., 2010).

A third possibility is that both of these mechanisms are at work. First, within-condition similarity may be lower in old adults than in young adults. Second, between-condition similarity may be higher in old adults than in young adults. Clearly, the operation of both of these mechanisms would also lead to reduced neural distinctiveness in old vs. young adults.

When examining these components individually, we found an age-related reduction in within-condition similarity but not in between-condition similarity. Thus, the greatest driver of age-related neural dedifferentiation in the present study was a decrease in the reliability of neural activation patterns (i.e., noisier neural activation patterns in the older adults) rather than increased representational dissimilarity for different categories per se. Consistent with this interpretation, Carp et al. (2011) also found that age-related changes in within-condition similarity were larger than age-related changes in between-condition similarity.

### 4.4 Higher GABA+ levels are associated with greater neural distinctiveness

Finally, we found that individual differences in GABA+ in ventral visual cortex were associated with individual differences in neural distinctiveness in the same region in older adults. Specifically, older adults with higher GABA+ levels tended to exhibit greater neural distinctiveness than those with lower GABA+ levels, even after controlling for gray matter volume and age. Notably, although visual GABA+ levels predicted neural distinctiveness in visual cortex, tissue-corrected sensorimotor and auditory GABA levels did not (although uncorrected auditory GABA levels did). These findings demonstrate a degree of regional-specificity in the relationship between GABA and neural distinctiveness.

The observed relationship between GABA and neural distinctiveness seems at least related to results from single cell studies conducted by Leventhal et al. (2003) who found that the application of a GABA agonist increased the selectivity of visual receptive fields in older macaques. Obviously, an fMRI measure of neural distinctiveness is very different from a single-unit measure of receptive field selectivity, but both findings suggest that GABA may be related to the specificity of neural representations.

Our results are also consistent with previous findings from the MiND study (Lalwani et al. 2019) that reported a similar relationship between GABA and neural distinctiveness, but in auditory cortex. The Lalwani et al. (2019) results were also region specific; GABA+ levels in visual cortex and sensorimotor cortex did not predict neural distinctiveness in auditory cortex. Furthermore, results from Cassady et al. (2019; 2020) observed relationships among sensorimotor neural distinctiveness, GABA+ levels, and network segregation. Together, these results suggest that regionally-specific relationships may exist between GABA levels and neural distinctiveness in the cortex of older adults.

One theoretical framework that seems consistent with our results is the revised Scaffolding Theory of Aging and Cognition (STAC-r) model (Reuter-Lorenz & Park, 2014). In this framework, brain structure (in our case, the GABAergic system) and brain function (in our case, neural distinctiveness) interact in a manner that subsequently influences cognitive performance during later adulthood. In particular, perhaps lower GABA+ levels reduce neural distinctiveness within ventral visual cortex within older adults, which then impairs cognitive processes such as fluid processing ability (Park et al., 2010).

Another possibility is that GABA+ levels are an indicator of neural reserve (Cabeza et al., 2018; Stern et al., 2018), such that older adults with higher GABA+ levels are better able to withstand other impairments in brain structure and function that are associated with aging. Longitudinal studies will assist in deepening our understanding of the processes by which GABA and neural distinctiveness interact in the context of cognitive decline during later adulthood.

### 4.5 Limitations

The current study has at least four limitations. 1) Like most imaging studies, the current study is correlational, and so we cannot conclude that reductions in ventral visual GABA+ levels cause neural dedifferentiation, just that the measures are associated. Additional research should examine longitudinal changes in GABA and neural distinctiveness to understand the coupling in the two measures as a lead-lag relationship would provide evidence for a possible causal role of GABA. 2) Another limitation of the current study (and other GABA MR spectroscopy studies in humans) is that the size of our spectroscopy voxels was large (3×3×3 cm; Gagnon et al., 2019). Our voxel placement therefore includes cortical tissue that probably is not directly involved in processing face and house information, which would presumably make it more difficult to detect a relationship between distinctiveness and GABA. 3) The fMRI data and MR spectroscopy data were collected during different scanning sessions on different days (Gagnon et al., 2019) so the coupling between them is not as tight as it could be. 4) Finally, MRS estimates of GABA do not measure GABA activity, but only GABA concentration. These limitations should presumably make it harder to observe relationships between GABA and distinctiveness, so our finding of a significant relationship suggests that the relationship may be fairly strong.

## 5.0 Conclusions

In closing, we report that GABA levels are reduced with age in ventral visual regions. We also found age-related neural dedifferentiation in older compared to younger adults in the same region. Finally, we demonstrated that individual differences in GABA levels were associated with individual differences in neural distinctiveness within older adults. These findings collectively are consistent with the hypothesis that age-related declines in GABA play a role in neural dedifferentiation within the ventral visual cortex.

## Acknowledgements

Funding was provided by a grant from the National Institute on Aging to TAP (R01AG050523)

## References

Abdulrahman, H., Fletcher, P. C., Bullmore, E., & Morcom, A. M. (2017). Dopamine and memory dedifferentiation in aging. NeuroImage, 153, 211–220. https://doi.org/10.1016/j.neuroimage.2015.03.031

Aufhaus, E., Weber-Fahr, W., Sack, M., Tunc-Skarka, N., Oberthuer, G., Hoerst, M., Meyer-Lindenberg, A., Boettcher, U., & Ende, G. (2013). Absence of changes in GABA concentrations with age and gender in the human anterior cingulate cortex: A MEGA-PRESS study with symmetric editing pulse frequencies for macromolecule suppression. Magnetic Resonance in Medicine, 69(2), 317–320. https://doi.org/10.1002/mrm.24257

Brooks, J. C. W., Beckmann, C. F., Miller, K. L., Wise, R. G., Porro, C. A., Tracey, I., & Jenkinson, M. (2008). Physiological noise modelling for spinal functional magnetic resonance imaging studies. NeuroImage, 39(2), 680–692. https://doi.org/10.1016/j.neuroimage.2007.09.018

Carp, J., Park, J., Hebrank, A., Park, D. C., & Polk, T. A. (2011). Age-related neural dedifferentiation in the motor system. PLOS ONE, 6(12), e29411. https://doi.org/10.1371/journal.pone.0029411

Carp, J., Park, J., Polk, T. A., & Park, D. C. (2011). Age differences in neural distinctiveness revealed by multi-voxel pattern analysis. NeuroImage, 56(2), 736–743. https://doi.org/10.1016/j.neuroimage.2010.04.267

Caspary, D. M., Milbrandt, J. C., & Helfert, R. H. (1995). Central auditory aging: GABA changes in the inferior colliculus. Experimental Gerontology, 30(3–4), 349–360.

Cassady, K., Gagnon, H., Freiburger, E., Lalwani, P., Simmonite, M., Park, D. C., Peltier, S. J., Taylor, S. F., Weissman, D. H., Seidler, R. D., & Polk, T. A. (2020). Network segregation varies with neural distinctiveness in sensorimotor cortex. NeuroImage, 212, 116663. https://doi.org/10.1016/j.neuroimage.2020.116663

Cassady, K., Gagnon, H., Lalwani, P., Simmonite, M., Foerster, B., Park, D., Peltier, S. J., Petrou, M., Taylor, S. F., Weissman, D. H., Seidler, R. D., & Polk, T. A. (2019). Sensorimotor network segregation declines with age and is linked to GABA and to sensorimotor performance. NeuroImage, 186, 234–244. https://doi.org/10.1016/j.neuroimage.2018.11.008

Edden, R. A. E., Puts, N. A. J., Harris, A. D., Barker, P. B., & Evans, C. J. (2014). Gannet: A batch-processing tool for the quantitative analysis of gamma-aminobutyric acid–edited MR spectroscopy spectra. Journal of Magnetic Resonance Imaging?: JMRI, 40(6), 1445–1452.

Gagnon, H., Simmonite, M., Cassady, K., Chamberlain, J. D., Freiburger, E., Lalwani, P., … Polk, T. A. (2018). Michigan Neural Distinctiveness (MiND) project: Investigating the scope, causes, and consequences of age-related neural dedifferentiation. bioRxiv, 466516. https://doi.org/10.1101/466516

Gao, F., Edden, R. A. E., Li, M., Puts, N. A. J., Wang, G., Liu, C., … Barker, P. B. (2013). Edited magnetic resonance spectroscopy detects an age-related decline in brain GABA levels. NeuroImage, 78, 75–82. https://doi.org/10.1016/j.neuroimage.2013.04.012

Gershon, R. C., Cella, D., Fox, N. A., Havlik, R. J., Hendrie, H. C., & Wagster, M. V. (2010). Assessment of neurological and behavioural function: the NIH Toolbox. The Lancet Neurology, 9(2), 138–139. https://doi.org/10.1016/S1474-4422(09)70335-7

Goh, J. O., Suzuki, A., & Park, D. C. (2010a). Reduced neural selectivity increases fMRI adaptation with age during face discrimination. NeuroImage, 51(1), 336–344. https://doi.org/10.1016/j.neuroimage.2010.01.107

Grachev, I. D., & Apkarian, A. V. (2001). Aging alters regional multichemical profile of the human brain: an in vivo1H-MRS study of young versus middle-aged subjects. Journal of Neurochemistry, 76(2), 582–593. https://doi.org/10.1046/j.1471-4159.2001.00026.x

Greene, H. A., & Madden, D. J. (1987). Adult age differences in visual acuity, stereopsis, and contrast sensitivity. American Journal of Optometry and Physiological Optics, 64(10), 749–753.

Greenhouse, I., Noah, S., Maddock, R. J., & Ivry, R. B. (2016). Individual differences in GABA content are reliable but are not uniform across the human cortex. NeuroImage, 139, 1–7. https://doi.org/10.1016/j.neuroimage.2016.06.007

Fandakova, Y., Leckey, S., Driver, C. C., Bunge, S. A., & Ghetti, S. (2019). Neural specificity of scene representations is related to memory performance in childhood. NeuroImage, 199, 105–113. https://doi.org/10.1016/j.neuroimage.2019.05.050

Harris, A. D., Puts, N. A. J., & Edden, R. A. E. (2015). Tissue correction for GABA-edited MRS: Considerations of voxel composition, tissue segmentation and tissue relaxations. Journal of Magnetic Resonance Imaging?: JMRI, 42(5), 1431–1440. https://doi.org/10.1002/jmri.24903

Haxby, J. V., Gobbini, M. I., Furey, M. L., Ishai, A., Schouten, J. L., & Pietrini, P. (2001). Distributed and overlapping representations of faces and objects in ventral temporal cortex. Science, 293(5539), 2425–2430. https://doi.org/10.1126/science.1063736

Haxby, J. V., Hoffman, E. A., & Gobbini, M. I. (2000). The distributed human neural system for face perception. Trends in Cognitive Sciences, 4(6), 223–233. https://doi.org/10.1016/S1364-6613(00)01482-0

Hermans, L., Leunissen, I., Pauwels, L., Cuypers, K., Peeters, R., Puts, N. A. J., Edden, R. A. E., & Swinnen, S. P. (2018). Brain GABA Levels Are Associated with Inhibitory Control Deficits in Older Adults. Journal of Neuroscience, 38(36), 7844–7851. https://doi.org/10.1523/JNEUROSCI.0760-18.2018

Hua, T., Kao, C., Sun, Q., Li, X., & Zhou, Y. (2008). Decreased proportion of GABA neurons accompanies age-related degradation of neuronal function in cat striate cortex. Brain Research Bulletin, 75(1), 119–125. https://doi.org/10.1016/j.brainresbull.2007.08.001

Isaacson, J. S., & Scanziani, M. (2011). How inhibition shapes cortical activity. Neuron, 72(2), 231–243. https://doi.org/10.1016/j.neuron.2011.09.027

Koen, J. D., Hauck, N., & Rugg, M. D. (2019). The relationship between age, neural differentiation, and memory performance. Journal of Neuroscience, 39(1), 149–162. https://doi.org/10.1523/JNEUROSCI.1498-18.2018

Koen, J. D., & Rugg, M. D. (2019). Neural Dedifferentiation in the Aging Brain. Trends in Cognitive Sciences, 23(7), 547–559. https://doi.org/10.1016/j.tics.2019.04.012

Lalwani P, Gagnon H, Cassady K, Simmonite M, Peltier SJ, Seidler RD, Taylor SF, Weissman DH, & Polk TA (2019). Neural distinctiveness declines with age in auditory cortex and is associated with auditory GABA levels. NeuroImage, 116033. https://doi.org/10.1016/j.neuroimage.2019.116033

Leventhal, A. G., Wang, Y., Pu, M., Zhou, Y., & Ma, Y. (2003). GABA and its agonists improved visual cortical function in senescent monkeys. Science, 300(5620), 812–815. https://doi.org/10.1126/science.1082874

Li, S.-C., & Lindenberger, U. (1999). Cross-level unification: A computational exploration of the link between deterioration of neurotransmitter systems and dedifferentiation of cognitive abilities in old age. In Cognitive neuroscience of memory (pp. 103–146). Ashland, OH, US: Hogrefe & Huber Publishers.

Li, S.-C., Lindenberger, U., & Sikström, S. (2001). Aging cognition: from neuromodulation to representation. Trends in Cognitive Sciences, 5(11), 479–486. https://doi.org/10.1016/S1364-6613(00)01769-1

Li, S.-C., & Sikström, S. (2002). Integrative neurocomputational perspectives on cognitive aging, neuromodulation, and representation. Neuroscience & Biobehavioral Reviews, 26(7), 795–808. https://doi.org/10.1016/S0149-7634(02)00066-0

Majdi M, Ribeiro-da-Silva A, & Cuello AC (2007). Cognitive impairment and transmitter-specific pre- and postsynaptic changes in the rat cerebral cortex during ageing. Eur. J.Neurosci. 26, 3583–3596.

Meng, M., Cherian, T., Singal, G., & Sinha, P. (2012). Lateralization of face processing in the human brain. Proceedings of the Royal Society B: Biological Sciences, 279(1735), 2052–2061. https://doi.org/10.1098/rspb.2011.1784

Miettinen R, Sirviö J, Riekkinen P, Laakso MP, & Riekkinen M (1993). Neocortical, hippocampal and septal parvalbumin- and somatostatin-containing neurons in young and aged rats: correlation with passive avoidance and water maze performance. Neuroscience 53, 367–378.

Nasreddine, Z. S., Phillips, N. A., Bédirian, V., Charbonneau, S., Whitehead, V., Collin, I., … Chertkow, H. (2005). The Montreal Cognitive Assessment, MoCA: a brief screening tool for mild cognitive impairment. Journal of the American Geriatrics Society, 53(4), 695–699. https://doi.org/10.1111/j.1532-5415.2005.53221.x

Park, D. C., Polk, T. A., Park, R., Minear, M., Savage, A., & Smith, M. R. (2004). Aging reduces neural specialization in ventral visual cortex. Proceedings of the National Academy of Sciences of the United States of America, 101(35), 13091–13095. https://doi.org/10.1073/pnas.0405148101

Park, J., Carp, J., Hebrank, A., Park, D. C., & Polk, T. A. (2010). Neural specificity predicts fluid processing ability in older adults. Journal of Neuroscience, 30(27), 9253–9259. https://doi.org/10.1523/JNEUROSCI.0853-10.2010

Park, J., Carp, J., Kennedy, K. M., Rodrigue, K. M., Bischof, G. N., Huang, C.-M., … Park, D. C. (2012). Neural broadening or neural attenuation? Investigating age-related dedifferentiation in the face network in a large lifespan sample. The Journal of Neuroscience?: The Official Journal of the Society for Neuroscience, 32(6), 2154–2158. https://doi.org/10.1523/JNEUROSCI.4494-11.2012

Pinto, J. G. A., Hornby, K. R., Jones, D. G., & Murphy, K. M. (2010). Developmental changes in GABAergic mechanisms in human visual cortex across the lifespan. Frontiers in Cellular Neuroscience, 4. https://doi.org/10.3389/fncel.2010.00016

Pitchaimuthu, K., Wu, Q., Carter, O., Nguyen, B. N., Ahn, S., Egan, G. F., & McKendrick, A. M. (2017). Occipital GABA levels in older adults and their relationship to visual perceptual suppression. Scientific Reports, 7(1). https://doi.org/10.1038/s41598-017-14577-5

Porges, E. C., Woods, A. J., Edden, R. A. E., Puts, N. A. J., Harris, A. D., Chen, H., … Cohen, R. A. (2017). Frontal gamma-aminobutyric acid concentrations are associated with cognitive performance in older adults. Biological Psychiatry?: Cognitive Neuroscience and Neuroimaging, 2(1), 38–44. https://doi.org/10.1016/j.bpsc.2016.06.004

R Core Team (2017) R: a language and environment for statistical computing. Vienna, Austria: R Foundation.

Schmolesky, M. T., Wang, Y., Pu, M., & Leventhal, A. G. (2000). Degradation of stimulus selectivity of visual cortical cells in senescent rhesus monkeys. Nature Neuroscience, 3(4), 384–390. https://doi.org/10.1038/73957

Simmonite, M., Carp, J., Foerster, B. R., Ossher, L., Petrou, M., Weissman, D. H., & Polk, T. A. (2018). Age-related declines in occipital GABA are associated with reduced fluid processing ability. Academic Radiology. https://doi.org/10.1016/j.acra.2018.07.024

Stanley, D. P., & Shetty, A. K. (2004). Aging in the rat hippocampus is associated with widespread reductions in the number of glutamate decarboxylase-67 positive interneurons but not interneuron degeneration. Journal of Neurochemistry, 89(1), 204–216. https://doi.org/10.1111/j.1471-4159.2004.02318.x

Stanley EM, Fadel JR, & Mott DD (2012). Interneuron loss reduces dendritic inhibition and GABA release in hippocampus of aged rats. Neurobiol. Aging 33, 431.e431–413.

St-Laurent, M., Abdi, H., Bondad, A., & Buchsbaum, B. R. (2014). Memory Reactivation in Healthy Aging: Evidence of Stimulus-Specific Dedifferentiation. Journal of Neuroscience, 34(12), 4175– 4186. https://doi.org/10.1523/JNEUROSCI.3054-13.2014

Voss, M. W., Erickson, K. I., Chaddock, L., Prakash, R. S., Colcombe, S. J., Morris, K. S., … Kramer, A. F. (2008). Dedifferentiation in the visual cortex: An fMRI investigation of individual differences in older adults. Brain Research, 1244, 121–131. https://doi.org/10.1016/j.brainres.2008.09.051

Zheng, L., Gao, Z., Xiao, X., Ye, Z., Chen, C., & Xue, G. (2018). Reduced fidelity of neural representation underlies episodic memory decline in normal aging. Cerebral Cortex, 28(7), 2283–2296. https://doi.org/10.1093/cercor/bhx130

