## Supplemental Materials for "GABA levels in ventral visual cortex decline with age and are associated with neural distinctiveness"

1) Raw GABA+ level age differences

Age had a significant effect on raw GABA+ estimates such that older adults exhibited reduced GABA+ levels compared to younger adults ($t(85.52) = 4.83$, $p < 0.001$). GABA+ levels were still reduced in older compared to younger adults after controlling for SNR differences (r(87 ) = -0.38, $p < .001$). The raw measurements in the left and right hemisphere voxels were significantly correlated (r(84) = .49, $p < .001$) and a mixed design ANOVA (age group as a between-subjects factor and hemisphere of voxel placement as a within-subjects factor) revealed no significant interaction between hemisphere and age ($F(1,84) = .25$, $p > .05$).

2) Raw GABA+ levels and neural distinctiveness

Older adults with higher levels of raw GABA+ exhibited increased neural distinctiveness ($\beta(49) = .27$, $p < 0.05$), even after controlling for individual differences in gray matter volume, age, and GABA+ levels within auditory and sensorimotor voxels ($\beta(45) = .40$, $p < 0.01$). Visual neural distinctiveness was most strongly related to visual GABA levels, although raw auditory GABA+ levels were also significantly related ($\beta(45) = -0.31$, $p < 0.05$). In contrast, sensorimotor GABA+ levels did not predict visual neural distinctiveness ($\beta(45) = -0.07$, $p > 0.05$).

3) Alternative functional ROI definition for neural distinctiveness estimation

At the recommendation of a reviewer, we defined our functional ROI following procedures conducted by Park et al., (2012). Specifically, we centered a circular ROI of 2,000 vertices on each participant's face and house peak within each hemisphere of the anatomical ROI. When defining our ROI in this manner, we again observed an age-related reduction in neural distinctiveness in older compared to younger adults ($t(62.77)=2.24$, $p < .05$), and ventral visual GABA+ levels were positively correlated with neural distinctiveness in older adults (r(49) = 0.31, $p < .05$). Ventral visual GABA+ also positively predicted neural distinctiveness when controlling for auditory GABA+, somatomotor GABA+ , age, and gray matter volume (ventral visual GABA+ $\beta(45) = 0.26$, $p < 0.05$; all other predictor *p*'s >.05).
